# DrugPattern: a web-based tool for drug set enrichment analysis

**DOI:** 10.1101/178632

**Authors:** Chuanbo Huang, Weili Yang, Junpei Wang, Yuan Zhou, Bin Geng, Georgios Kararigas, Jichun Yang, Qinghua Cui

## Abstract

Set enrichment analysis based methods (e.g. gene set enrichment analysis) have provided great helps in mining patterns in biomedical datasets, however, tools for inferring regular patterns in drug-related datasets are still limited. For the above purpose, here we developed a web-based tool, DrugPattern. DrugPattern first collected and curated 7019 drug sets, including indications, adverse reaction, targets, pathways etc. For a list of interested drugs, DrugPattern then evaluates the significance of the enrichment of these drugs in each of the 7019 drug sets. To validate DrugPattern, we applied it to predict the potential protective roles of oxidized low-density lipoprotein (oxLDL), a widely accepted deleterious factor for the body. We predicted that oxLDL has beneficial effects on some diseases, most of which were supported by literature except type 2 diabetes (T2D), in which oxLDL was previously believed to be a risk factor. Animal experiments further validated that oxLDL indeed has beneficial effects on T2D. These data confirmed the prediction accuracy of our approach and revealed unexpected protective roles for oxLDL in various diseases including T2D. This study provides a tool to infer regular patterns in biomedical datasets based on drug set enrichment analysis.

## INTRODUCTION

Recently, the performances of techniques in data collection, storage, processing, and analysis have changed dramatically(1). As a result, the amount of data in both biology and medicine has increased remarkably(2). So far, the open-access big data presents us unprecedented opportunities for scientific discovery in health and medicine(2). Big data analyses (BDA) has recently been used to discover and predict, for example, novel biomarkers of cardiovascular disease(3), trends of flu(4), new cancer targets(5), and novel subtypes of type 2 diabetes(6). These studies indicate that BDA is showing great potential both to infer knowledge (7) and to predict unappreciated information for guiding personalized and precision medicine(8). Among the biomedical BDA tools, set enrichment analysis based methods play very important roles in mining rules and patterns, for example gene set enrichment analysis tools (e.g. DAVID Bioinformatics(9) and GSEA(10)), miRNA set enrichment analysis tools (e.g. TAM(11) and miEAA(12)), metabolite set enrichment analysis (e.g. MSEA(13) and MPEA(14)), and phenotype set enrichment analysis (e.g. PSEA(15)). Recently, as the fast development of techniques such as drug-repurposing(16) and network pharmacology(17), many studies often generate a list of drugs. However, tools to discover regular patterns in a list of drugs based on drug set enrichment analysis are still limited. Recently, Napolitano et al. developed a “drug-set enrichment analysis” tool, DSEA(18), to investigate drug mode of action. However, DSEA is not doing enrichment analysis for drug sets but for gene set and gene pathway to the list of genes whose expression profiles induced by one drug. Therefore, a tool directly doing enrichment analysis for drug sets is needed.

Accumulating datasets in public drug databases provides us an opportunity to collect and curate a variety of drug sets. We define a drug set as a list of drugs that are grouped by the same or similar rules, for example the drugs with the same targets could be a drug set. Here we first collected and curated 7019 drug sets from public databases. This makes it possible to infer enriched patterns in a new drug list. For doing so, here we developed a drug set enrichment analysis tool, DrugPattern. To confirm the accuracy of DrugPattern, we applied it to predict the potential beneficial effects of oxidized low-density lipoprotein (oxLDL). LDL, especially the oxidized one, is referred to bad cholesterol and its lowering therapy has been an important strategy for the protection and prevention of coronary heart disease but faced with unexpected puzzles in the new century(19). To apply DrugPattern to oxLDL, we need to get a list of drugs that have similar biological effects with oxLDL. For doing so, we identified the drugs that have highly similar induced gene profiles with oxLDL. Then we performed enrichment analysis of these drugs in the drug sets for Disease category. The results showed that oxLDL was predicted to have beneficial effects on a number of diseases, including bacterial infection, type 2 diabetes (T2D), cancer, and depression. Literature mining found evidence to support most of the above predictions but T2D is an exception. Currently, oxLDL was known as a risk factor on T2D. Thus, animal experiments had been performed to validate whether oxLDL is beneficial or deleterious to T2D. The results showed that oxLDL treatment significantly reduced levels of the fasting blood glucose, serum insulin, triglyceride and cholesterol, and improved the overall glucose intolerance and the global insulin sensitivity as well, suggesting that oxLDL has beneficial effects on T2D. These results confirmed the predictions of DrugPattern.

## MATERIALS AND METHODS

### Gene expression datasets and analysis

We downloaded processed gene expression profiles before and after oxLDL treatment of human ARPE-19 cells (retinal pigmented epithelial cells) from the GEO database (<u>https://www.ncbi.nlm.nih.gov/geo/,</u> GEO accession number: GSE5741). We then calculated the gene expression fold change after oxLDL treatment. We downloaded the gene expression fold change of ∼1000 drugs from the CMap database(20). Next, we searched the CMap drugs that are highly similar with oxLDL in their gene profile fold changes with the genes regulated by oxLDL (Rho>=0.06 & p-value<=1.0e-16, Spearman’s correlation). As a result, we got 66 oxLDL-similar drugs in gene-expression signatures (Supplementary Table 1).

### Enrichment analysis of oxLDL-similar drugs

We first downloaded drug-related data from databases such as DrugBank(21), KEGG (http://www.kegg.jp/), and ADReCS(22). We then processed these data into various drug sets, which referred to groups of drugs with the same biomedical item. For example, drugs with the same molecular target are assigned into one drug set. By this way, we collected 7019 drug sets. Next, we developed a web-based tool, DrugPattern (<u>http://www.cuilab.cn/drugpattern</u>), to perform drug set enrichment analysis for a drug list (e.g. the 66 oxLDL-similar drugs) in drug sets based on hypergeometric test. Assuming that P is the number of drugs included in all drug sets, S is the number of drugs included in drug set A, HP is the number of input drugs included in P, and HS represents the number of drugs that are of interest included in S. The probability of HS drugs of interest in drug set A is shown as Formula (1).

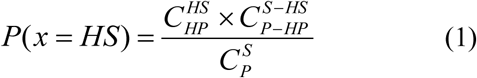

Here “ C ” means the combination operation. Therefore, the statistical significance of the enrichment of drugs of interest in drug set A is represented by Formula (2).

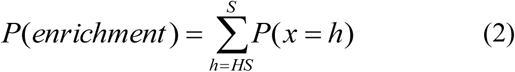

Finally, the p-values were corrected by FDR as well.

### Experimental mice

Male 8-10 week old C57BL/6J mice (weight 18-22 g) were employed in this study. All animal care and experimental protocols complied with the Animal Management Rules of the Ministry of Health of the People’s Republic of China and the guide for the Care and Use of the Laboratory Animals of the Peking University. All animal protocols were approved by the Animal Research Committee of the Peking University Health Science Center.

### Administration of HFD-induced diabetic mice with human oxLDL

Mice were fed a 45% HFD for 16 weeks before experimental assays. The HFD-fed mice were randomly divided into two groups based on OGTT. Human oxLDL was purchased from Dalian Meilun Biology Technology Co. LTD (China) and dissolved in sterile PBS. The experimental group of mice were daily treated with human oxidized LDL at the dose of 1.5 mg/kg bodyweight via the tail vein injection as previously described elsewhere (23,24) for 3 weeks, whereas the control group of mice were treated with equal volume of sterile PBS. The fasting blood glucose levels were monitored weekly. On the14^th^ and 18^th^ day post oxLDL injection, OGTT and ITT were performed respectively. On the 22^th^ day, mice were sacrificed. Serum and tissues were collected for further experimental assays.

### Metabolic phenotyping

OGTT was performed in 6 hours (8am to 2pm) fasted mice using a dose of 3g/kg glucose, and blood glucose levels were monitored at 0, 15, 30, 60, 90 and 120 minutes post glucose load using a Freestyle brand glucometer (Roche) via blood collected from the tail(25). The blood glucose concentrations at 0 minute were determined as fasting blood glucose. Plasma insulin levels were measured using the Rat/Mouse Insulin ELASA kit (MILIPORE) (26-28) at fed states before sacrifice. ITT using a dose of 1 U/kg insulin was performed in 4 h fasted mice (10am to 2pm). Tail vein blood was used for measuring the glucose concentration with a glucometer (Roche) at 0, 15, 30, 60, 90 and 120 minutes post insulin load.

### Measuring the hepatic and serum triglyceride and cholesterol content

The hepatic lipid content were determined as previously described (26-28). In brief, certain weight of liver tissue (20-40mg) was homogenized in 1ml chloroform:methanol (2:1 vol/vol) solvent (Beijing Chemical Technology) on ice. The homogenized samples were incubated for 16 hours at 4°C, and then 300 μl distilled water was added into each sample. The samples were mixed by vortexing, and then centrifuged at 12,000 rpm for 10 min at 4°C. The clear organic phase was collected and dried off under a stream of nitrogen. The dried pellets were finally solubilized in PBS buffer containing 5% Triton X-100. The total triglycerides (TG) and total cholesterol (TC) were quantified by using TG and CHO assay kits (BIOSINO Bio-Technology and Science Inc.). Serum triglyceride (TG) and cholesterol levels were also determined with these assay kits according to the manufacturer’s instructions.

### Real-Time PCR Assay

Liver and other tissues were quickly dissected from mice and snap frozen in liquid nitrogen. Total RNA was extracted using RNApure High-purity Total RNA Rapid Extraction Kit (Spin-column) (BioTeke Corporation, Beijing, China) according to the manufacturer’s recommendations. After DNase digestion, total RNA was eluted with 30∼100 µl of RNase-free water and stored at -80°C. Purity and quantity of extracted total RNA was determined by the ratio of absorbance at A260 to A280, and agarose gel electrophoresis. For analysis of mRNA expression, reverse transcription was performed with the use of RevertAidTM First Strand cDNA Synthesis Kit (Fermentas K1622) according to the manufacturer’s protocol using oligo (dT) primers. Real-time PCR experiments wer performed on Real-Time qPCR System (Agilent Technologies, Stratagene Mx3000P). The relative fold change in mRNA expression was calculated using the 2-ΔΔ*C*t method. All primer sequences for quantitative PCR assays are listed in Supplementary Table 2.

### Statistical analysis

Data are presented as mean ± S.E.M. Statistical significance of differences between groups was analyzed by unpaired Student’s *t* test or by one-way analysis of variance (ANOVA) when more than two groups were compared.

## RESULTS AND DISCUSSION

### The construction of DrugPattern and the application of DrugPattern to oxLDL

We collected and curated 7019 drug sets, which cover several big categories, such as target, pathway, chemical classification, disease, and adverse reaction etc. We then developed a web-based tool, DrugPattern (<u>http://www.cuilab.cn/drugpattern</u>), to discover regular patterns in a given drug list. To validate DrugPattern, we focus on oxLDL. As shown in **Figure 1a**, we first dissected the gene-expression profiles induced by oxLDL treatment (**Materials and methods**). Next, by searching the gene-expression profiles induced by more than 1000 drugs in the Connectivity Map (CMap) database, which was originally designed for drug-repurposing, we obtained the drugs that show high similarity in gene-expression profiles with oxLDL. As a result, we got 66 oxLDL-similar drugs in gene-expression profiles (see **Materials and Methods**, Supplementary Table 1). Then, we enter these drugs into DrugPattern for enrichment analysis of these drugs in the 7019 drug sets. Finally, we predicted the beneficial effects of oxLDL (that is, the enriched disease-related drug sets of the oxLDL-similar drugs, **Figure 1b**).

**Figure 1.**
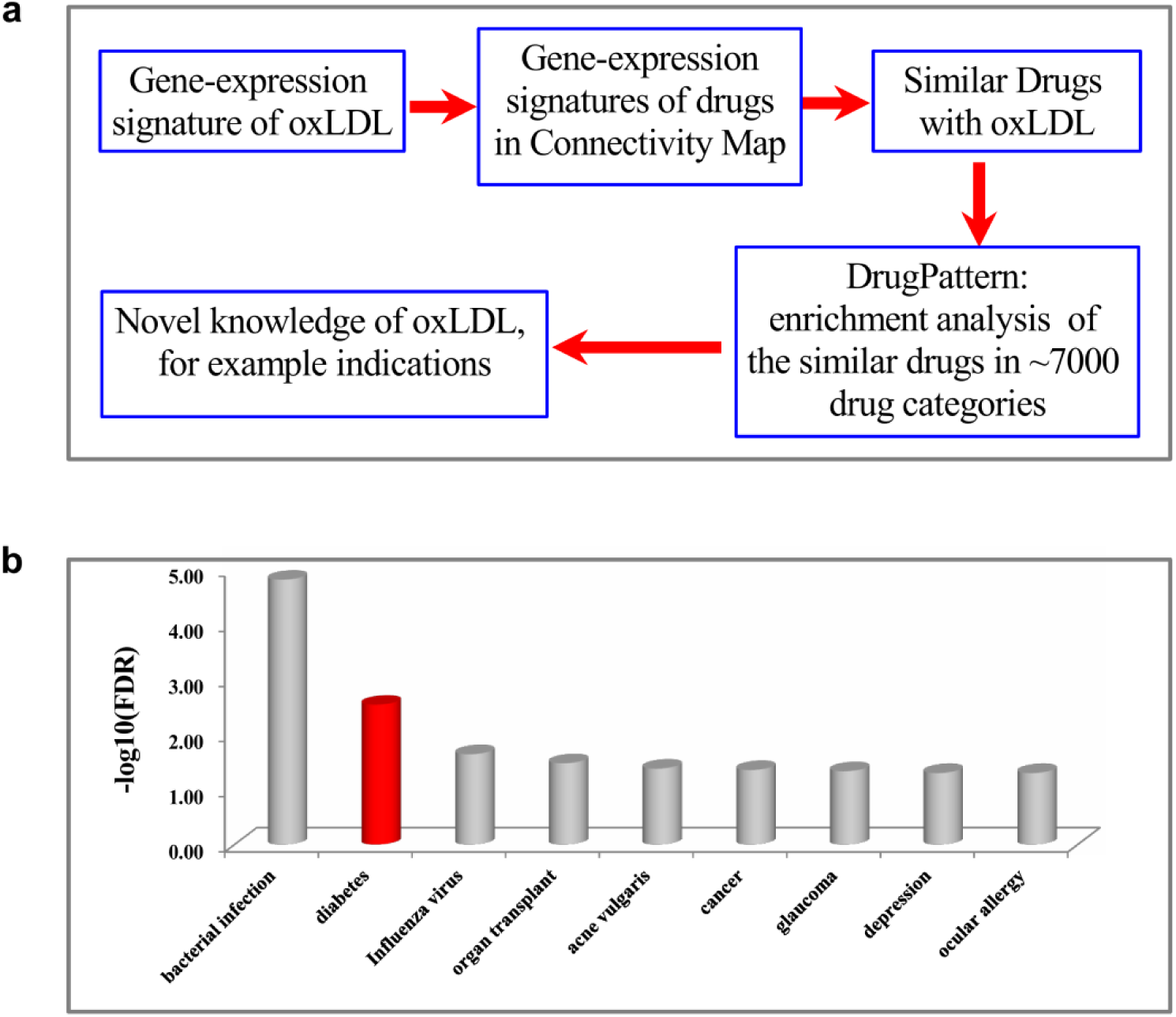
An *in silico* framework for the inference of causality and degrees of causality based on drug-related big data using oxLDL as an example. **(a)** and the prediction results for potentially beneficial effects of oxLDL **(b).** The length of each bar represents the degree of causality. The prediction for diabetes (the red bar) was selected to confirm experimentally.

The link of LDL, a cholesterol-carrying particle, to coronary heart disease as a major risk factor represents one of the greatest biomedical stories in the 20^th^ century(19). Moreover, LDL-lowering therapy has been an important strategy for the protection and prevention of coronary heart disease(19). However, in the new century, accumulating studies have provided unexpected observations, for example, the association between total cholesterol (TC) and all-cause mortality is negative but not positive(29,30). Given that the association between TC and cardiovascular disease is positive(31), cholesterol should have protective roles in other diseases. Therefore, it is becoming timely necessary to evaluate the roles of LDL, the cholesterol carrier, which is critical for personalized and precision medicine, for example, to find the people who should receive LDL-lowering and the people who cannot receive LDL-lowering. Here we then used the above tool to infer the beneficial effects for oxLDL, which is considered more deleterious than LDL. As a result, we predicted that oxLDL has beneficial effects on a number of diseases (**Figure 1b**), including bacterial infection, type 2 diabetes, leprostatic, influenza virus, cancer, glaucoma, and depression. Literature mining further supported that high cholesterol may indeed protect against infection(32), cancer(30), and depression(33), suggesting that DrugPattern has a good prediction accuracy. For the prediction of diabetes, higher-level oxLDL was widely observed in type 2 diabetes patients(34), which triggers some researchers to hypothesize that it has harmful effects on type 2 diabetes(34). However, it still lacks clear evidence the relationship between oxLDL and type 2 diabetes is beneficial, harmful, or just passive.

### Animal experiments confirmed the beneficial effects of oxLDL on type 2 diabetes

Given the beneficial effects of oxLDL on type 2 diabetes predicted by DrugPattern, here we further performed animal experiments to validate this prediction. For doing so, mice fed on high-fat diet (HFD) for 4 months were treated with human oxLDL for 3 weeks. In clinical practice, obese type 2 diabetic patients are always suggested to control their bodyweight and energy intake. To mimic these clinical strategies, HFD was replaced with normal diet once the mice began receiving oxLDL treatment. During the period of oxLDL administration, the bodyweight of both groups of mice decreased gradually, but there was no difference between them at each time point (Figure 2a). OxLDL treatment also had no effect on the weight of liver, white adipose, heat and kidney (data not shown). At 7, 14 and 21 days post oxLDL treatment, the fasting blood glucose of oxLDL-treated mice was significantly lower than that of control mice (Fig. 2b). Oral glucose tolerance test revealed that the overall glucose intolerance was significantly improved at 14 days post oxLDL treatment (Fig. 2c-f). Insulin tolerance test indicated that the global insulin sensitivity was also improved at 18 days post oxLDL treatment (Fig. 3a-b). In support, serum insulin level was decreased after oxLDL treatment (Fig. 3c). Both serum TG and CHO levels were also significantly reduced after oxLDL treatment (Fig. 3d-e). OxLDL treatment slightly increased serum AST and ALT activities, but not statistically different when compared with control mice (Fig. 4a-b). Gluconeogenic PEPCK and G6Pase mRNAs were reduced, whereas that of FAS, SREBP-1, FAM3A and FAM3B remained unchanged in oxLDL-treated mouse livers when compared with control mouse livers (Fig. 5a-c). This suggested the suppression of hepatic gluconeogenesis, further supporting the amelioration of fasting hyperglycemia after oxLDL treatment. However, it should be noted that oxLDL administration had no significant beneficial effect on hyperglycemia and global insulin resistance on HFD-induced diabetic mice when they continued receiving HFD during time of oxLDL treatment (Supplemental Fig. 1).

**Figure 2.**
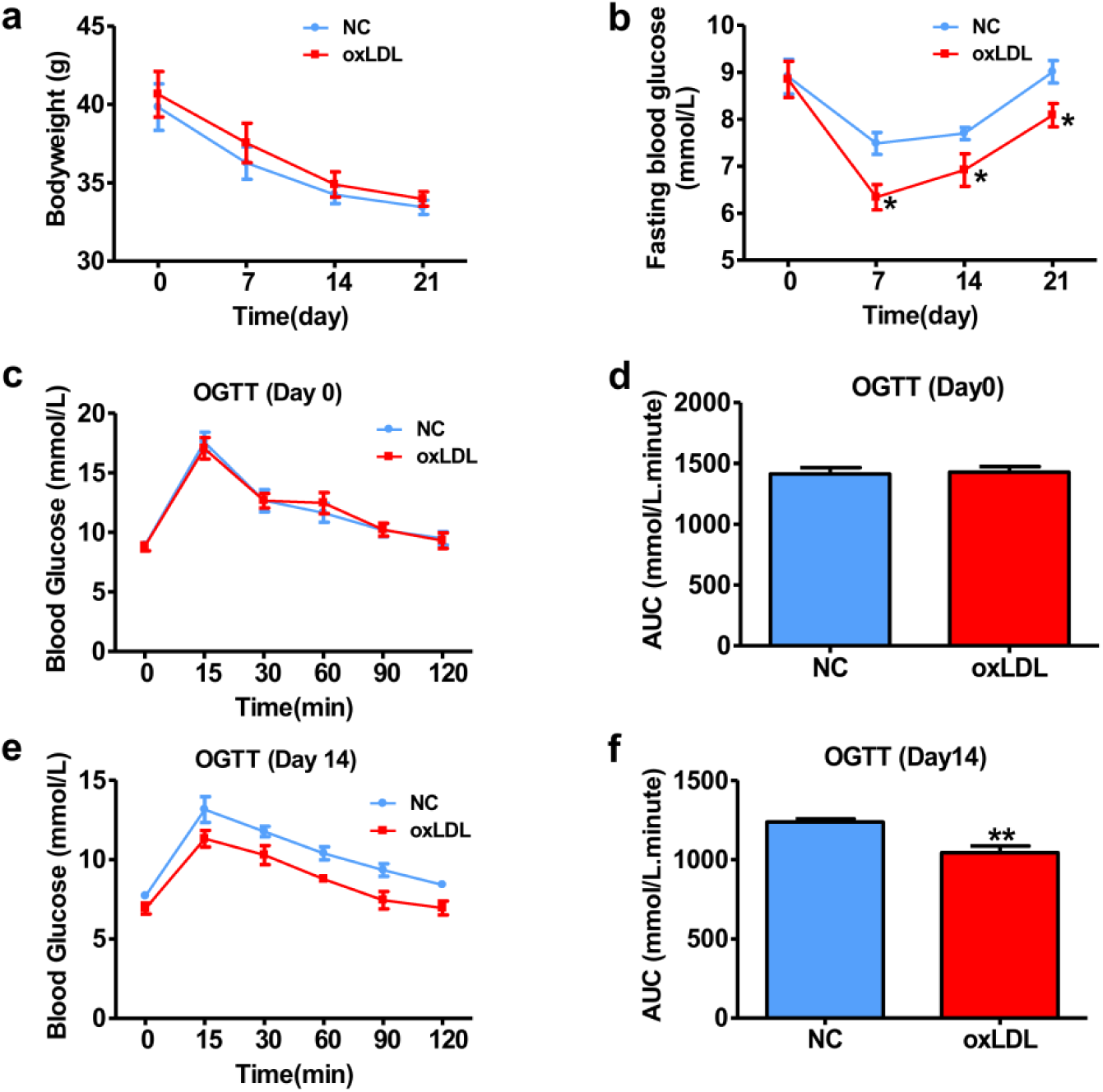
oxLDL treatment improved hyperglycemia of HFD mice. Mice were fed on HFD (high fat diet) for 4 months, and then treated with oxLDL or placebo (PBS buffer) for 3 weeks. Upon treatment onset, the diet was replaced with normal diet (ND). (**a**) Body weight. (**b**) Fasting blood glucose levels. (**c-d**) OGTT prior treatment onset and AUC data, respectively. (**e-f**) OGTT 14 days post treatment onset and AUC data. respectively. The blood glucose at 0 minute was also presented as fasting blood glucose in panel b. NC, control mice treated with placebo (PBS); oxLDL, mice treated with oxLDL. N=9, *P<0.05, **P<0.01 compared with control mice.

**Figure 3.**
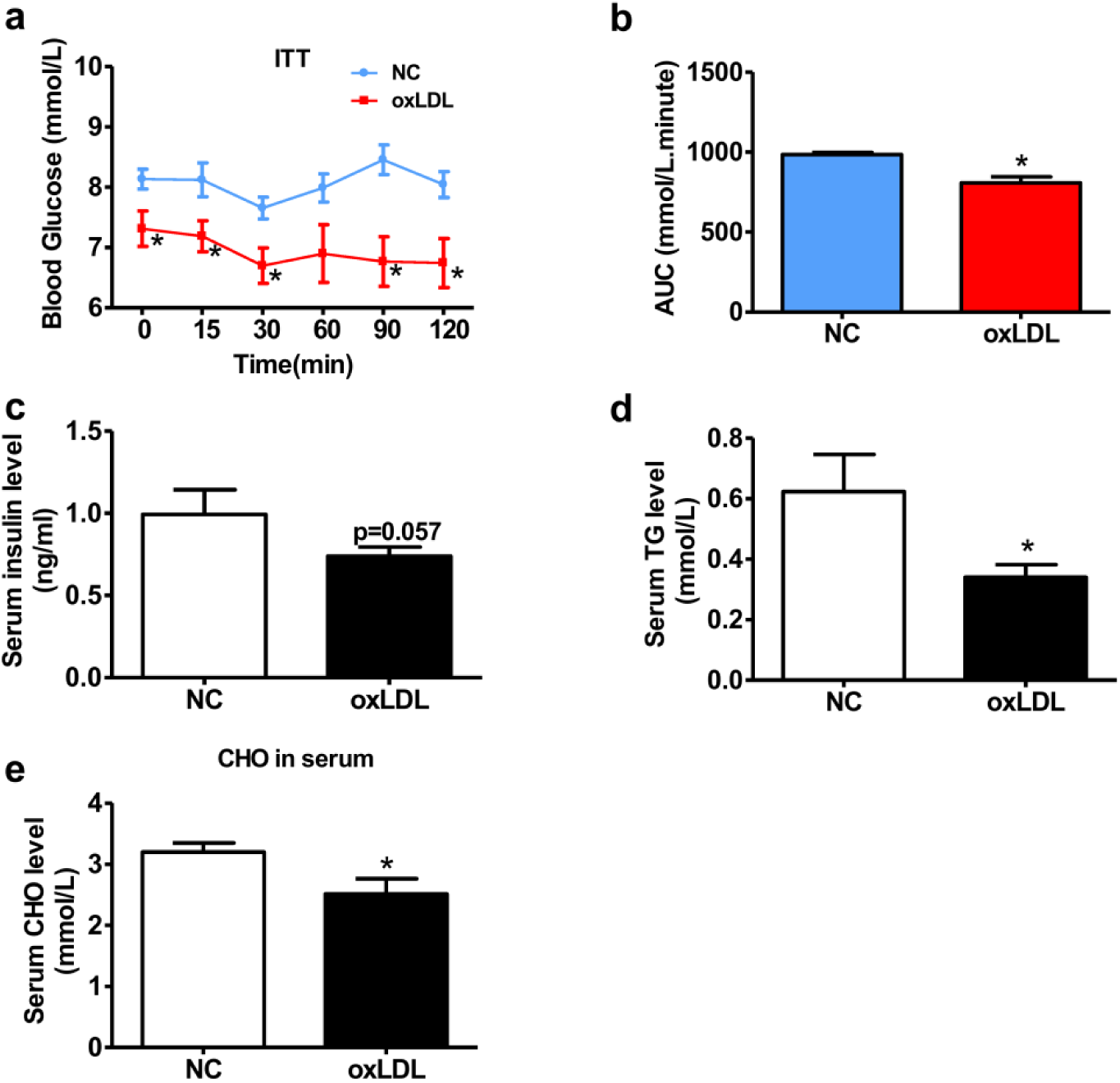
oxLDL treatment increased global insulin sensitivity of HFD mice. (**a-b**) ITT of mice at 18 days post treatment onset and AUC data, respectively. (**c**) Serum insulin level in HFD mice treated with oxLDL or PBS. Mice were sacrificed at 22 days post treatment onset. (**d-e**) OxLDL treatment reduced serum TG (**d**) and CHO (**e**) levels in HFD mice. TG, triglycerides; CHO, cholesterol. N=9, *P<0.05, **P<0.01 compared with control mice.

**Figure 4.**
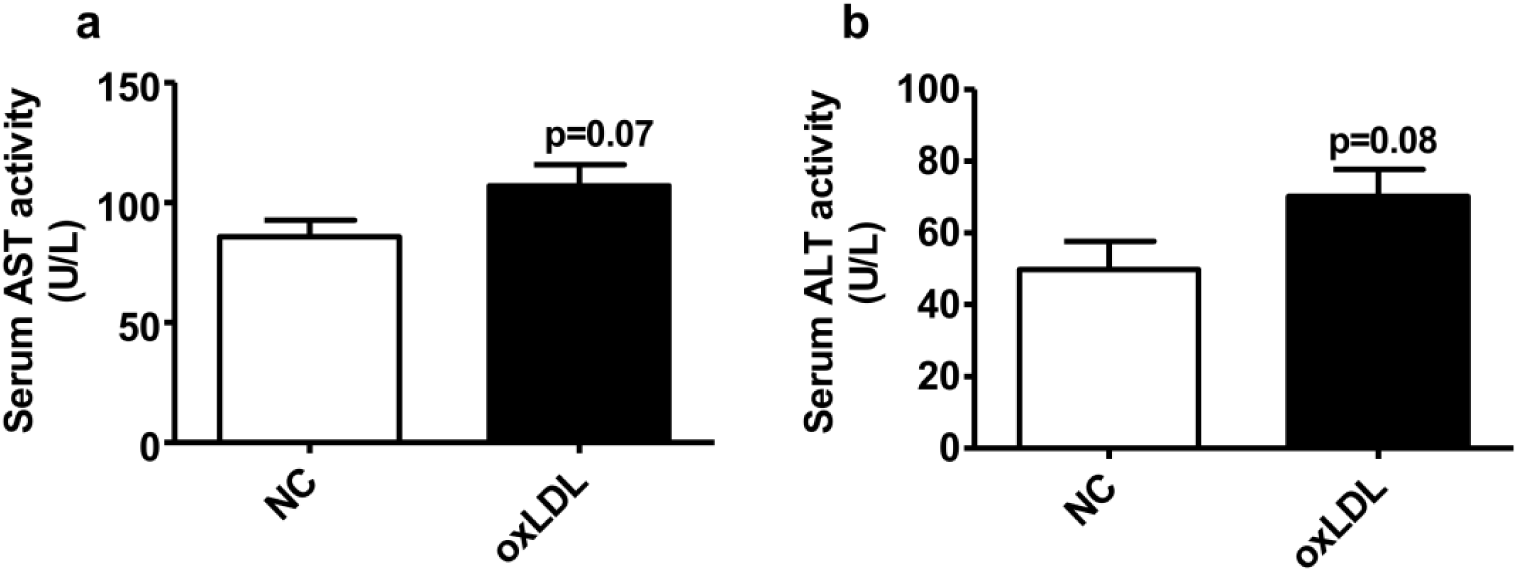
Serum AST and ALT activities. (**a-b**) The activities of serum AST and ALT were measured in oxLDL-treated and control mice. AST, aspartate transaminase; ALT, alanine aminotransferase. N=9.

**Figure 5.**
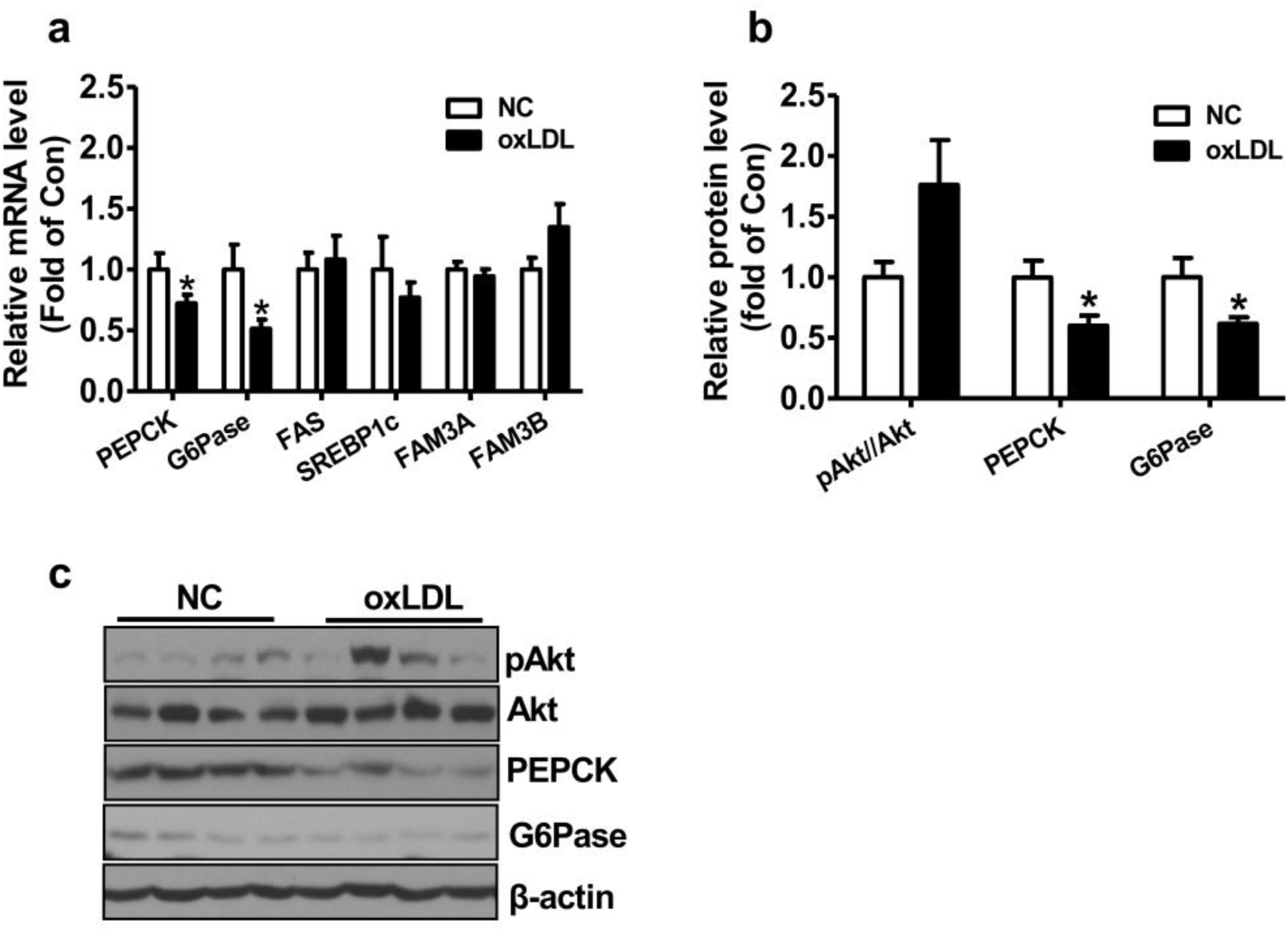
Gluconeogenic factor levels in mouse livers. (**a**) The mRNA levels of glucose and lipid metabolizing genes in mouse livers after treatment onset. (**b-c**) The levels of gluconeogenic protein in mouse livers after treatment onset. Representative immunoblotting images are shown in panel c and quantitative data in panel b. N=9, *P<0.05 compared with control mice.

### Discussion

In this study, we have developed a web-based tool, DrugPattern, to discover regular patterns in a list of drugs. DrugPattern collected and curated 7019 drug sets, which cover a number of big categories, such as drug targets, pathways, chemical classification, diseases, and adverse reaction etc. For a given list of drugs, DrugPattern then calculated the enrichment of these drugs in each of the 7019 drug sets. We confirmed the usefulness of DrugPattern by using it to predict the beneficial effects (the enriched disease-related drug sets) of oxLDL. As a result, we predicted that oxLDL may have beneficial effects on infection, type 2 diabetes, depression and cancer etc. OxLDL had been recognized as an oxidative marker in human and animals, and is a risk factor of various diseases(35,36). However, each molecule may have many faces in different cellular processes in various organ or diseases. For example, restrict of cholesterol intake had been long believed to have beneficial effects on human health. However, Dietary Guidelines Advisory Committee (DGAC) had cancelled the restrict on daily cholesterol intake in <US Dietary *Guideline 2015>* based on the clinical trials in the past decades(37). In support, one recent systematic review revealed that high LDL-cholesterol is negatively correlated with mortality in people over 60 years(30). This finding had challenged the current cholesterol hypothesis, in which it is widely believed that high LDL-cholesterol is deleterious for the body. DrugPattern predicted that oxLDL may exert beneficial effects in various diseases including cancer, diabetes and depression. In support of our prediction, it had been reported that serum cholesterol level is inversely correlated with cancer(38), and some drugs that reduce serum lipids including LDL and oxLDL will increase the prevalence of cancer(39). Moreover, another systematic review had revealed that low serum LDL level is associated with depression(33). Beyond our prediction, oxLDL also inhibits hepatitis C virus (HCV) infection (40,41). OxLDL also upregulates miR-146a to inhibit macrophage activation in cultured human THP-1 macrophages (42). Clearly, oxLDL does exert beneficial effects in some diseases, at least in certain stage of the disease progression. Although circulating LDL and oxLDL levels were increased in type 2 diabetic patients(43,44) and reduced after control of hyperglycemia(45), their precise roles in the pathogenesis of diabetes still remains unclear. The possibility that an increase in serum LDL and subsequent oxLDL levels is a protective response to hyperglycemia, at least in certain stage of diabetes, can not be precluded. Statins are widely used as lipid-lowering drugs that reduce LDL and oxLDL in human and animals(46,47). However, it had been reported that statins will increase the prevalence of diabetes(48-50). One more recent meta-analysis report further revealed that LDL-C-lowering genetic variants were associated with a higher risk of type 2 diabetes(51). These important findings had revealed that low serum LDL-cholesterol is associated with high risk of diabetes. In the current study, we further provided direct experimental evidence that administration of oxLDL improved fasting hyperglycemia, glucose intolerance and insulin resistance in a HFD-fed diabetic mouse model restricting energy intake. Collectively, the experimental results and literature evidence support the predictions by DrugPattern, suggested that the roles of oxLDL in some diseases such as type 2 diabetes may not be as what we had thought, at least in certain stage of disease progression. The availability of this tool allows for fast scientific discovery and will facilitate the investigation of big data based precision medicine.

## ACKNOWLEDGEMENTS

We thank all members of Prof. Cui’s and Prof. Yang’s groups for useful discussions.

## REFERENCES

1. Mayer-Schonberger, V. (2016) Big Data for cardiology: novel discovery? Eur Heart J, 37, 996–1001.

2. Rumsfeld, J.S., Joynt, K.E. and Maddox, T.M. (2016) Big data analytics to improve cardiovascular care: promise and challenges. Nat Rev Cardiol, 13, 350–359.

3. Gerstein, H.C., Pare, G., McQueen, M.J., Haenel, H., Lee, S.F., Pogue, J., Maggioni, A.P., Yusuf, S., Hess, S. and Outcome Reduction With Initial Glargine Intervention Trial, I. (2015) Identifying Novel Biomarkers for Cardiovascular Events or Death in People With Dysglycemia. Circulation, 132, 2297–2304.

4. Ginsberg, J., Mohebbi, M.H., Patel, R.S., Brammer, L., Smolinski, M.S. and Brilliant, L. (2009) Detecting influenza epidemics using search engine query data. Nature, 457, 1012–1014.

5. Jiang, P. and Liu, X.S. (2015) Big data mining yields novel insights on cancer. Nat Genet, 47, 103–104.

6. Li, L., Cheng, W.Y., Glicksberg, B.S., Gottesman, O., Tamler, R., Chen, R., Bottinger, E.P. and Dudley, J.T. (2015) Identification of type 2 diabetes subgroups through topological analysis of patient similarity. Sci Transl Med, 7, 311ra174.

7. Chaussabel, D. and Pulendran, B. (2015) A vision and a prescription for big data-enabled medicine. Nat Immunol, 16, 435–439.

8. de Lemos, J.A., Rohatgi, A. and Ayers, C.R. (2015) Applying a Big Data Approach to Biomarker Discovery: Running Before We Walk? Circulation, 132, 2289–2292.

9. Huang da, W., Sherman, B.T. and Lempicki, R.A. (2009) Systematic and integrative analysis of large gene lists using DAVID bioinformatics resources. Nature protocols, 4, 44–57.

10. Subramanian, A., Tamayo, P., Mootha, V.K., Mukherjee, S., Ebert, B.L., Gillette, M.A., Paulovich, A., Pomeroy, S.L., Golub, T.R., Lander, E.S. et al. (2005) Gene set enrichment analysis: a knowledge-based approach for interpreting genome-wide expression profiles. Proceedings of the National Academy of Sciences of the United States of America, 102, 15545–15550.

11. Lu, M., Shi, B., Wang, J., Cao, Q. and Cui, Q. (2010) TAM: a method for enrichment and depletion analysis of a microRNA category in a list of microRNAs. BMC bioinformatics, 11, 419.

12. Backes, C., Khaleeq, Q.T., Meese, E. and Keller, A. (2016) miEAA: microRNA enrichment analysis and annotation. Nucleic acids research, 44, W110–116.

13. Xia, J. and Wishart, D.S. (2010) MSEA: a web-based tool to identify biologically meaningful patterns in quantitative metabolomic data. Nucleic acids research, 38, W71–77.

14. Kankainen, M., Gopalacharyulu, P., Holm, L. and Oresic, M. (2011) MPEA-metabolite pathway enrichment analysis. Bioinformatics, 27, 1878–1879.

15. Ried, J.S., Doring, A., Oexle, K., Meisinger, C., Winkelmann, J., Klopp, N., Meitinger, T., Peters, A., Suhre, K., Wichmann, H.E. et al. (2012) PSEA: Phenotype Set Enrichment Analysis-a new method for analysis of multiple phenotypes. Genetic epidemiology, 36, 244–252.

16. Sleire, L., Forde-Tislevoll, H.E., Netland, I.A., Leiss, L., Skeie, B.S. and Enger, P.O. (2017) Drug repurposing in cancer. Pharmacological research, 124, 74–91.

17. Boezio, B., Audouze, K., Ducrot, P. and Taboureau, O. (2017) Network-based Approaches in Pharmacology. Molecular informatics.

18. Napolitano, F., Sirci, F., Carrella, D. and di Bernardo, D. (2016) Drug-set enrichment analysis: a novel tool to investigate drug mode of action. Bioinformatics, 32, 235–241.

19. Goldstein, J.L. and Brown, M.S. (2015) A century of cholesterol and coronaries: from plaques to genes to statins. Cell, 161, 161–172.

20. Lamb, J., Crawford, E.D., Peck, D., Modell, J.W., Blat, I.C., Wrobel, M.J., Lerner, J., Brunet, J.P., Subramanian, A., Ross, K.N. et al. (2006) The Connectivity Map: using gene-expression signatures to connect small molecules, genes, and disease. Science, 313, 1929–1935.

21. Law, V., Knox, C., Djoumbou, Y., Jewison, T., Guo, A.C., Liu, Y., Maciejewski, A., Arndt, D., Wilson, M., Neveu, V. et al. (2014) DrugBank 4.0: shedding new light on drug metabolism. Nucleic acids research, 42, D1091–1097.

22. Cai, M.C., Xu, Q., Pan, Y.J., Pan, W., Ji, N., Li, Y.B., Jin, H.J., Liu, K. and Ji, Z.L. (2015) ADReCS: an ontology database for aiding standardization and hierarchical classification of adverse drug reaction terms. Nucleic acids research, 43, D907–913.

23. Calara, F., Dimayuga, P., Niemann, A., Thyberg, J., Diczfalusy, U., Witztum, J.L., Palinski, W., Shah, P.K., Cercek, B., Nilsson, J. et al. (1998) An animal model to study local oxidation of LDL and its biological effects in the arterial wall. Arterioscler Thromb Vasc Biol, 18, 884–893.

24. Wang, J.J., Chen, X.L., Xu, C.B., Jiang, G.F., Lin, J., Liu, E.Q., Qin, X.P. and Li, J. (2016) The ERK1/2 pathway participates in the upregulation of the expression of mesenteric artery alpha1 receptors by intravenous tail injections of mmLDL in mice. Vascul Pharmacol, 77, 80–88.

25. Yang, Z., Wang, X., Wen, J., Ye, Z., Li, Q., He, M., Lu, B., Ling, C., Wu, S. and Hu, R. (2011) Prevalence of non-alcoholic fatty liver disease and its relation to hypoadiponectinaemia in the middle-aged and elderly Chinese population. Arch Med Sci, 7, 665–672.

26. Li, J., Chi, Y., Wang, C., Wu, J., Yang, H., Zhang, D., Zhu, Y., Wang, N., Yang, J. and Guan, Y. (2011) Pancreatic-derived factor promotes lipogenesis in the mouse liver: role of the Forkhead box 1 signaling pathway. Hepatology, 53, 1906–1916.

27. Wang, C., Chen, Z., Li, S., Zhang, Y., Jia, S., Li, J., Chi, Y., Miao, Y., Guan, Y. and Yang, J. (2014) Hepatic overexpression of ATP synthase beta subunit activates PI3K/Akt pathway to ameliorate hyperglycemia of diabetic mice. Diabetes, 63, 947–959.

28. Wang, C., Chi, Y., Li, J., Miao, Y., Li, S., Su, W., Jia, S., Chen, Z., Du, S., Zhang, X. et al. (2014) FAM3A activates PI3K p110alpha/Akt signaling to ameliorate hepatic gluconeogenesis and lipogenesis. Hepatology, 59, 1779–1790.

29. Hamazaki, T., Okuyama, H., Ogushi, Y. and Hama, R. (2015) Towards a Paradigm Shift in Cholesterol Treatment. A Re-examination of the Cholesterol Issue in Japan. Ann Nutr Metab, 66 Suppl 4, 1–116.

30. Ravnskov, U., Diamond, D.M., Hama, R., Hamazaki, T., Hammarskjold, B., Hynes, N., Kendrick, M., Langsjoen, P.H., Malhotra, A., Mascitelli, L. et al. (2016) Lack of an association or an inverse association between low-density-lipoprotein cholesterol and mortality in the elderly: a systematic review. BMJ Open, 6, e010401.

31. Prospective Studies, C., Lewington, S., Whitlock, G., Clarke, R., Sherliker, P., Emberson, J., Halsey, J., Qizilbash, N., Peto, R. and Collins, R. (2007) Blood cholesterol and vascular mortality by age, sex, and blood pressure: a meta-analysis of individual data from 61 prospective studies with 55,000 vascular deaths. Lancet, 370, 1829–1839.

32. Ravnskov, U. (2003) High cholesterol may protect against infections and atherosclerosis. QJM, 96, 927–934.

33. Persons, J.E. and Fiedorowicz, J.G. (2016) Depression and serum low-density lipoprotein: A systematic review and meta-analysis. J Affect Disord, 206, 55–67.

34. Marin, M.T., Dasari, P.S., Tryggestad, J.B., Aston, C.E., Teague, A.M. and Short, K.R. (2015) Oxidized HDL and LDL in adolescents with type 2 diabetes compared to normal weight and obese peers. J Diabetes Complications, 29, 679–685.

35. Steinberg, D. (1997) Lewis A. Conner Memorial Lecture. Oxidative modification of LDL and atherogenesis. Circulation, 95, 1062–1071.

36. Gradinaru, D., Borsa, C., Ionescu, C. and Prada, G.I. (2015) Oxidized LDL and NO synthesis-Biomarkers of endothelial dysfunction and ageing. Mech Ageing Dev, 151, 101–113.

37. Meeting, D. (December 15, 2014).

38. Ravnskov, U., McCully, K.S. and Rosch, P.J. (2012) The statin-low cholesterol-cancer conundrum. QJM, 105, 383–388.

39. Newman, T.B. and Hulley, S.B. (1996) Carcinogenicity of lipid-lowering drugs. JAMA, 275, 55–60.

40. Westhaus, S., Bankwitz, D., Ernst, S., Rohrmann, K., Wappler, I., Agne, C., Luchtefeld, M., Schieffer, B., Sarrazin, C., Manns, M.P. et al. (2013) Characterization of the inhibition of hepatitis C virus entry by in vitro-generated and patient-derived oxidized low-density lipoprotein. Hepatology, 57, 1716–1724.

41. von Hahn, T., Lindenbach, B.D., Boullier, A., Quehenberger, O., Paulson, M., Rice, C.M. and McKeating, J.A. (2006) Oxidized low-density lipoprotein inhibits hepatitis C virus cell entry in human hepatoma cells. Hepatology, 43, 932–942.

42. Li, Z., Wang, S., Zhao, W., Sun, Z., Yan, H. and Zhu, J. (2016) Oxidized low-density lipoprotein upregulates microRNA-146a via JNK and NF-kappaB signaling. Mol Med Rep, 13, 1709–1716.

43. Bahadoran, Z., Mirmiran, P., Hosseinpanah, F., Rajab, A., Asghari, G. and Azizi, F. (2012) Broccoli sprouts powder could improve serum triglyceride and oxidized LDL/LDL-cholesterol ratio in type 2 diabetic patients: a randomized double-blind placebo-controlled clinical trial. Diabetes Res Clin Pract, 96, 348–354.

44. Hoogeveen, R.C., Ballantyne, C.M., Bang, H., Heiss, G., Duncan, B.B., Folsom, A.R. and Pankow, J.S. (2007) Circulating oxidised low-density lipoprotein and intercellular adhesion molecule-1 and risk of type 2 diabetes mellitus: the Atherosclerosis Risk in Communities Study. Diabetologia, 50, 36–42.

45. Galland, F., Duvillard, L., Petit, J.M., Lagrost, L., Vaillant, G., Brun, J.M., Gambert, P. and Verges, B. (2006) Effect of insulin treatment on plasma oxidized LDL/LDL-cholesterol ratio in type 2 diabetic patients. Diabetes Metab, 32, 625–631.

46. Hofnagel, O., Luechtenborg, B., Weissen-Plenz, G. and Robenek, H. (2007) Statins and foam cell formation: impact on LDL oxidation and uptake of oxidized lipoproteins via scavenger receptors. Biochim Biophys Acta, 1771, 1117–1124.

47. Mikhailidis, D.P. and Athyros, V.G. (2014) Dyslipidaemia in 2013: New statin guidelines and promising novel therapeutics. Nat Rev Cardiol, 11, 72–74.

48. Culver, A.L., Ockene, I.S., Balasubramanian, R., Olendzki, B.C., Sepavich, D.M., Wactawski-Wende, J., Manson, J.E., Qiao, Y., Liu, S., Merriam, P.A. et al. (2012) Statin use and risk of diabetes mellitus in postmenopausal women in the Women's Health Initiative. Arch Intern Med, 172, 144–152.

49. Ma, Y., Culver, A., Rossouw, J., Olendzki, B., Merriam, P., Lian, B. and Ockene, I. (2013) Statin therapy and the risk for diabetes among adult women: do the benefits outweigh the risk? Ther Adv Cardiovasc Dis, 7, 41–44.

50. Ridker, P.M., Pradhan, A., MacFadyen, J.G., Libby, P. and Glynn, R.J. (2012) Cardiovascular benefits and diabetes risks of statin therapy in primary prevention: an analysis from the JUPITER trial. Lancet, 380, 565–571.

51. Lotta, L.A., Sharp, S.J., Burgess, S., Perry, J.R., Stewart, I.D., Willems, S.M., Luan, J., Ardanaz, E., Arriola, L., Balkau, B. et al. (2016) Association Between Low-Density Lipoprotein Cholesterol-Lowering Genetic Variants and Risk of Type 2 Diabetes: A Meta-analysis. JAMA, 316, 1383–1391.

